# Canine *Leishmania* infection in communities with notified human visceral leishmaniasis cases in Albania

**DOI:** 10.64898/2026.11.25.769359

**Authors:** Teita Myrseli, Kujtim Mersini, Yaarit Nachum-Biala, Silva Bino, Gad Baneth

## Abstract

Visceral leishmaniasis (VL) is an important zoonosis in Albania, where dogs are the main reservoir of *Leishmania infantum*. This retrospective study describes *Leishmania* infection in dogs sampled in 2015 in communities where human VL cases had been notified. After 16 human cases were notified in the first half of 2015, household and neighbouring dogs were sampled in June and July 2015 in 19 locations across 11 districts. Sera from 305 dogs were tested by an inhouse ELISA, and 178 whole blood samples by kinetoplast DNA realtime PCR, followed by internal transcribed spacer 1 (ITS1) high resolution melt PCR and sequencing. Ten dogs (3.3%; 95% confidence interval: (1.6 - 5.9) were seropositive, and 22 (7.2%) had borderline results. Parasite DNA was detected in 5 of 178 dogs (2.8%), and ITS1 sequences identical to *L. infantum* reference sequences were obtained from two dogs. None of the six dogs living in case households was seropositive. Because sampling was targeted, these proportions describe the sampled dogs only. The results document canine infection in these communities in 2015 and provide a baseline for future surveillance; a population-based survey is needed to assess the current situation.

**Simple Summary:** Visceral leishmaniasis is a severe disease of people caused by a microscopic parasite that sand flies transmit, and dogs are the parasite’s main animal reservoir in the Mediterranean region. In 2015, sixteen human cases were notified in Albania. Veterinary teams then sampled 305 dogs living in the same villages and city neighbourhoods as the patients. Ten dogs carried antibodies against the parasite, and its genetic material was detected in the blood of five of the 178 dogs tested, so dogs in these communities were infected at the time. Positive dogs were found in seven of the eleven districts investigated, but none of the six dogs living in the patients’ own households was positive for antibodies. These findings document infection in dogs around human cases during a period when national information on the disease was scarce, and they provide a baseline for future surveys. Monitoring dogs in areas where human cases occur would help detect and control the disease earlier.

## Introduction

Albania is endemic for leishmaniasis, and visceral leishmaniasis (VL) remains an important public health problem, with sustained transmission across multiple districts (Bino et al., 2016; Myrseli et al., 2011, 2014, 2021). Between 2005 and 2020, Albania had the highest cumulative incidence of autochthonous VL in Europe, although the reported incidence declined over that period (Maia et al., 2023). Human leishmaniasis is also re-emerging in the wider Balkan region (Vaselek, 2021), and VL notifications have increased in neighbouring Greece (Tzani et al., 2021). The main aetiological agent of leishmaniasis in Albania is *Leishmania infantum* (zymodeme MON-1), which has been documented in human cases, dogs and vectors (Cani, Petrela, et al., 2001; Velo et al., 2017). Transmission occurs through the bite of phlebotomine sand flies; *L. infantum* DNA has been detected in sand flies in Albania, and *Phlebotomus neglectus* has been incriminated as the principal vector species in the country (Ayhan et al., 2016; Velo et al., 2017).

Dogs are the main domestic reservoir of *L. infantum* and play a central role in maintaining transmission to humans (EFSA Panel on Animal Health and Welfare [EFSA AHAW Panel], 2015). Although leishmaniosis in dogs was recorded in Albania as early as 1937 (Food Safety and Veterinary Institute archive, file no. 4, unpublished), evidence has largely relied on isolated reports and limited surveys. The first parasitological confirmation of canine leishmaniosis in Albania was reported in 1977 (Adhami & Murati, 1977), and subsequent reports included parasite isolation and characterisation of *L. infantum* MON-1 (Cani, Petrela, et al., 2001; Miho et al., 1989). In the late 1990s, canine infection was increasingly observed in additional areas, reflecting changes in dog population dynamics and movement (Kero & Xinxo, 1998). Later cross-sectional surveys reported variable seroprevalence in selected districts, ranging from low positivity in Tirana, Durrës and Kamëz to substantially higher estimates in some foci (Lazri et al., 2008; Myrseli et al., 2009). Most infected dogs remain clinically healthy for long periods yet can infect sand flies, which makes the canine reservoir difficult to detect and control (Morales-Yuste et al., 2022; Travi et al., 2018).

Despite these observations, Albania still lacks a systematic national canine surveillance programme, and, to our knowledge, no data have been published on *Leishmania* infection in dogs living in the households and communities of notified human VL cases. In neighbouring countries, recent canine serosurveys have been carried out in Greece (Symeonidou et al., 2021), and *L. infantum* has been detected for the first time in sand flies in Kosovo (Xhekaj et al., 2023). Climatic suitability for *L. infantum* transmission has increased across Europe over the past two decades (Carvalho et al., 2024), and canine serosurveys triggered by notified human cases have recently been used elsewhere in Europe (Laidoudi et al., 2024). The aim of this retrospective study was to describe the frequency and geographical distribution of *Leishmania* infection in dogs sampled in 2015 in communities where human VL cases had been notified, and thereby to provide a documented baseline for future surveillance.

## Materials and methods

During the first half of 2015, 16 human VL cases were reported to the Institute of Public Health, Tirana. The cases had been notified through the national surveillance system for infectious diseases. A suspected case was defined as a person living in or having travelled to an endemic area who had fever lasting more than one week and clinical signs compatible with VL, or in whom there was a strong laboratory-based suspicion of the disease. A confirmed case was defined as such a person with clinical signs of VL and serological and/or parasitological confirmation. Field activities were conducted in areas linked to these notifications. Nineteen locations in 11 districts (Dibër, Elbasan, Fier, Gjirokastër, Kruja, Kukës, Kurbin, Lezha, Lushnja, Saranda and Tirana) were selected on the basis of information available from the case notifications, including place of residence and reported travel history, and of the presence of domestic dogs in the respective locations. Sampling was thus triggered by the human case notifications; the study was not designed to assess an epidemiological association between individual human cases and infected dogs.

Sampling included dogs from households in which human VL cases had been reported and dogs residing in the same communities but not belonging to those households. Sampling was carried out in June and July 2015 in collaboration with the regional veterinary authorities. For each dog, breed, primary use (guard, shepherd, hunting, pet or stray), setting (urban or rural) and terrain (mountainous, hilly or plain) were recorded. Dogs were examined at sampling, and clinical signs compatible with canine leishmaniosis (dermatological lesions, alopecia, weight loss and ocular abnormalities) were recorded. Blood was collected by cephalic or jugular venepuncture into vacutainer tubes without anticoagulant (for serum) and with EDTA (for whole blood). Serum was separated and stored until analysis, and whole blood was retained for molecular testing.

Anti-*Leishmania* antibodies were detected in 305 serum samples using an in-house ELISA based on *L. infantum* antigen, as described previously (Baneth et al., 1998). Results were classified as positive, borderline or negative; borderline results were not counted as positive. Molecular testing was performed on 178 whole-blood samples using real-time PCR targeting kinetoplast DNA (kDNA) with primers LEISH-1/LEISH-2 and the corresponding TaqMan probe, following a published protocol (Francino et al., 2006). Samples positive by kDNA real-time PCR were further analysed by internal transcribed spacer 1 (ITS1) high-resolution melt (HRM) PCR with primers ITS1-F-219F/ITS1-R-219R (Talmi-Frank et al., 2010).

ITS1 amplicons were sequenced and compared with sequences in GenBank using the Basic Local Alignment Search Tool (BLAST). The ITS1 sequences and the GenBank comparison are provided in Supplementary File 1.

The proportions of seropositive and PCR-positive dogs were calculated with exact (Clopper–Pearson) binomial 95% confidence intervals (CI). Because sampling was targeted rather than population-based, these proportions describe the sampled dogs only and are not estimates of prevalence in the district or national canine populations. Given the small number of positive dogs, results by district, setting, terrain and primary use are presented descriptively, without formal hypothesis testing. The spatial distribution of positive dogs and human cases was mapped using a geographic information system (GIS).

## Results

A total of 305 dogs from 19 locations in 11 districts were included in the serological survey. The sampled dogs included household and community animals from urban and rural settings. Approximately 80% were local mixed-breed dogs, and the remainder were pure-bred or crossbred dogs of other breeds. By primary use, 221 dogs were guard dogs, 35 shepherd dogs, 23 hunting dogs, 19 pet dogs and 7 stray dogs; the stray category is too small to be regarded as representative of the free-roaming dog population. Among the 16 households with notified human VL cases, six owned dogs, all of which were sampled; none of these dogs was seropositive by ELISA.

Overall, 10 of 305 dogs were seropositive (3.3%; 95% CI: 1.6–5.9), and a further 22 dogs (7.2%; 95% CI: 4.6–10.7) had borderline results. If borderline results were counted as positive, the proportion would be 32 of 305 (10.5%; 95% CI: 7.3–14.5). District-level results are shown in Table I. Seropositive dogs were found in six of the 11 districts, and eight of the ten were from rural settings. The locations of seropositive dogs and notified human VL cases in 2015 are shown in Figure 1.

**Table I.**
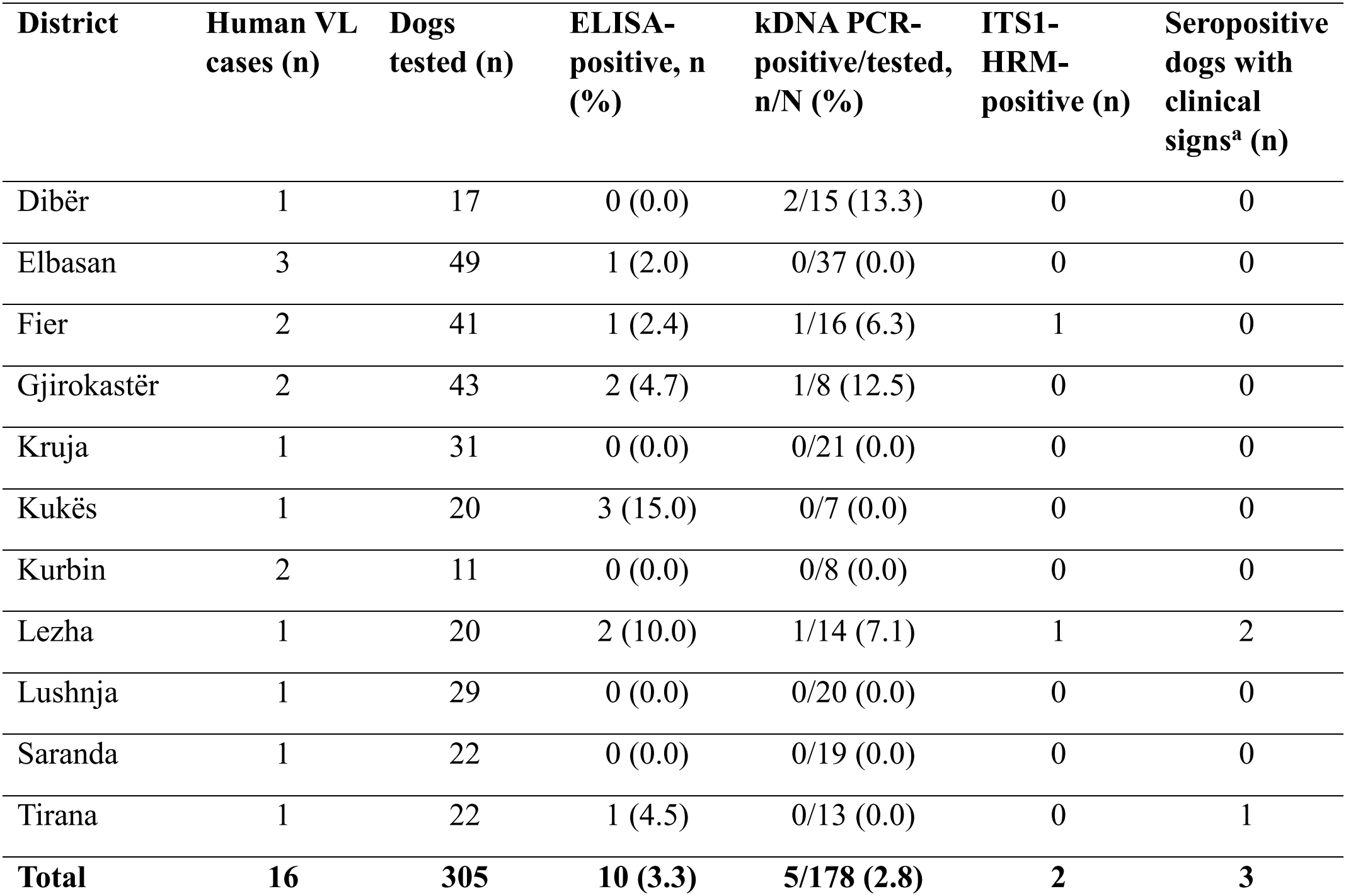
Notified human visceral leishmaniasis cases, dogs tested, and dogs positive by ELISA, kDNA real-time PCR and ITS1 high-resolution melt PCR, by district, Albania, 2015. ELISA = enzyme-linked immunosorbent assay; kDNA = kinetoplast DNA; PCR = polymerase chain reaction; ITS1-HRM = internal transcribed spacer 1 high-resolution melt PCR, performed on kDNA-positive samples only. Percentages are given in parentheses. ^a^Clinical signs compatible with canine leishmaniosis, recorded in seropositive dogs.

| **District** | **Human VL cases (n)** | **Dogs tested (n)** | **ELISA-positive, n (%)** | **kDNA PCR-positive/tested, n/N (%)** | **ITS1-HRM-positive (n)** | **Seropositive dogs with clinical signs<sup>a</sup> (n)** |
|---|---|---|---|---|---|---|
| Dibër | 1 | 17 | 0 (0.0) | 2/15 (13.3) | 0 | 0 |
| Elbasan | 3 | 49 | 1 (2.0) | 0/37 (0.0) | 0 | 0 |
| Fier | 2 | 41 | 1 (2.4) | 1/16 (6.3) | 1 | 0 |
| Gjirokastër | 2 | 43 | 2 (4.7) | 1/8 (12.5) | 0 | 0 |
| Kruja | 1 | 31 | 0 (0.0) | 0/21 (0.0) | 0 | 0 |
| Kukës | 1 | 20 | 3 (15.0) | 0/7 (0.0) | 0 | 0 |
| Kurbin | 2 | 11 | 0 (0.0) | 0/8 (0.0) | 0 | 0 |
| Lezha | 1 | 20 | 2 (10.0) | 1/14 (7.1) | 1 | 2 |
| Lushnja | 1 | 29 | 0 (0.0) | 0/20 (0.0) | 0 | 0 |
| Saranda | 1 | 22 | 0 (0.0) | 0/19 (0.0) | 0 | 0 |
| Tirana | 1 | 22 | 1 (4.5) | 0/13 (0.0) | 0 | 1 |
| **Total** | **16** | **305** | **10 (3.3)** | **5/178 (2.8)** | **2** | **3** |

**Figure 1.**
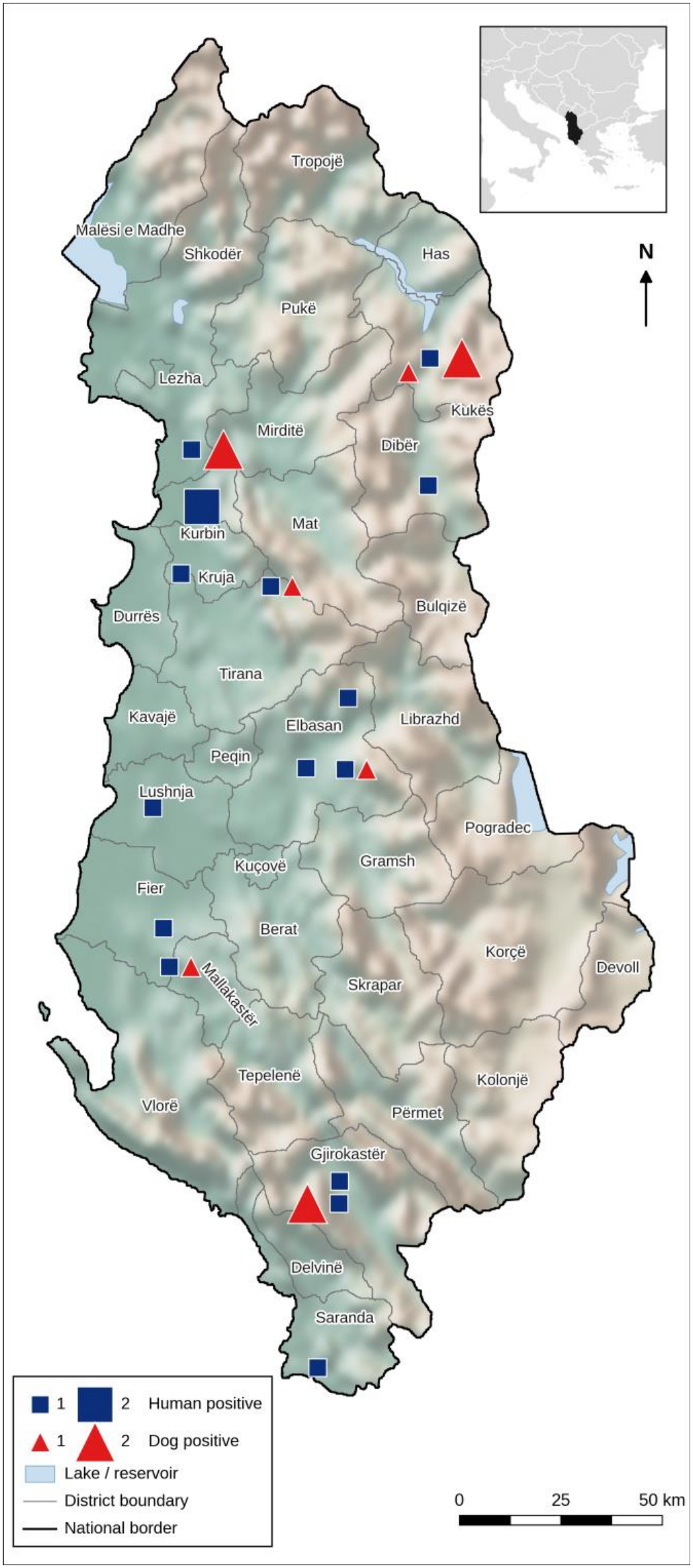
Locations of seropositive dogs and notified human visceral leishmaniasis cases, by district, Albania, 2015 (situation at the time of sampling).

Nine of the ten seropositive dogs were local mixed-breed dogs. The distribution of seropositive dogs by primary use is shown in Table II; the numbers in each category are too small for comparison.

**Table II.** Dogs seropositive for anti-*Leishmania* antibodies (ELISA) by primary use, Albania, 2015. CI = exact (Clopper–Pearson) binomial confidence interval. Proportions describe the sampled dogs only; the categories are too small for comparison, and the stray category (n = 7) is not representative of the free-roaming dog population.

| **Primary use** | **Dogs tested (n)** | **ELISA-positive, n (%)** | **95% CI (%)** |
|---|---|---|---|
| Guard | 221 | 8 (3.6) | 1.6–7.0 |
| Shepherd | 35 | 1 (2.9) | 0.1–14.9 |
| Hunting | 23 | 1 (4.3) | 0.1–21.9 |
| Pet | 19 | 0 (0.0) | 0.0–17.6 |
| Stray | 7 | 0 (0.0) | 0.0–41.0 |
| **Total** | **305** | **10 (3.3)** | **1.6–5.9** |

Clinical signs compatible with canine leishmaniosis, including dermatological lesions, alopecia, weight loss and ocular abnormalities, were recorded in three seropositive dogs (two in Lezha and one in Tirana; Table I). The remaining seven seropositive dogs showed no clinical signs at the time of examination.

Parasite DNA was detected by kDNA real-time PCR in 5 of 178 dogs tested (2.8%; 95% CI: 0.9–6.4). The proportion of dogs tested by PCR varied by district from 19% in Gjirokastër (8/43) to 88% in Dibër (15/17) (Table I), so PCR results cannot be compared between districts. The five PCR-positive dogs were from Dibër (two), Fier, Gjirokastër and Lezha. Both PCR-positive dogs from Dibër were ELISA-negative or borderline, as no dog in this district was seropositive. ITS1 HRM PCR was positive in two of the five kDNA-positive dogs, from Fier and Lezha. The two ITS1 sequences (218 bp) were identical to each other and showed 100% query coverage and 100% identity with *L. infantum* ITS1 sequences in GenBank (e.g. accession numbers KU975157.1, KU975158.1 and KU975159.1; Supplementary File 1). The same fragment is, however, also identical to some GenBank sequences attributed to *Leishmania donovani* and therefore does not discriminate between the two species on its own.

## Discussion

To our knowledge, this study provides the first data on canine infection from the households and communities of notified human VL cases in Albania. In this retrospective survey of dogs sampled in 2015 in communities where human VL cases had been notified, 10 of 305 dogs (3.3%) were seropositive and parasite DNA was detected in 5 of 178 dogs tested (2.8%). The detection of PCR-positive dogs demonstrates infection with *Leishmania* among dogs in the sampled communities at the time of sampling; the data cannot show whether transmission persisted, nor do they describe the current situation. Species-level identification was obtained for two dogs only. Their ITS1 sequences were identical to *L. infantum* reference sequences, but because the fragment is also shared by some sequences attributed to *L. donovani*, attribution to *L. infantum* rests on the epidemiological context: *L. infantum* is the only member of the *L. donovani* complex reported in Albania (Cani, Petrela, et al., 2001; Velo et al., 2017). The low proportion of dogs with clinical signs is consistent with the clinical spectrum of canine leishmaniosis, in which many infected dogs are asymptomatic yet can infect sand flies (Baneth et al., 2008; EFSA AHAW Panel, 2015). These data document canine infection in a period for which national human VL data were incomplete in international reporting (Maia et al., 2023).

The proportion of seropositive dogs observed here cannot be compared directly with earlier Albanian surveys (Cani, Myrseli, et al., 2001; Cicko et al., 1999; Myrseli et al., 2009) or with recent surveys elsewhere in the Mediterranean region (Laidoudi et al., 2024; Symeonidou et al., 2021). These studies differ in sampling frame, diagnostic assay and cut-off, dog population and epidemiological setting; in the present study, sampling was targeted at communities with notified human cases rather than being population-based. The large borderline group illustrates how strongly such proportions depend on the cut-off: counting borderline results as positive would raise the proportion from 3.3% to 10.5%. Borderline reactions may reflect recent or low-level infection, past exposure or non-specific reactivity. Serology and PCR also reflect different stages of infection: antibodies can persist beyond detectable parasitaemia, whereas PCR sensitivity varies with the tissue sampled and the parasite burden (Francino et al., 2006). Similar discordance between serology and PCR has been reported in asymptomatic dogs in southern France (Hidé et al., 2024). That not all kDNA-positive dogs were positive by ITS1 PCR is probably explained by the higher sensitivity of kDNA PCR compared with ITS1 PCR, as reported previously (Baneth et al., 2020).

Sampling was triggered by human VL notifications, but the study does not demonstrate an epidemiological association between individual human cases and infected dogs. None of the six dogs living in case households was seropositive. This does not exclude a role of dogs in local transmission, which may involve dogs outside the index households (EFSA AHAW Panel, 2015), but the present data cannot address this question. Gaps in surveillance also affect human data: Albania reported no human VL cases to the World Health Organization for 2013–2015 (Maia et al., 2023), although 16 cases were notified nationally in the first half of 2015 alone.

This study has limitations. The samples were collected in 2015, and the results describe the situation at that time; they should not be interpreted as estimates of the current frequency of canine infection, and no comparable recent data are available from the same communities. Only 178 of the 305 dogs were tested by PCR, and the proportion tested varied between districts, which may have introduced selection bias. The molecular data are limited to five PCR-positive dogs and two short ITS1 sequences, and the number of positive dogs was too small for comparison between groups. Finally, the age and sex of the dogs, the use of preventive measures against sand flies and the travel history of the dogs were not included in the analysis, so local acquisition of infection cannot be assumed for every dog.

## Conclusion

In 2015, *Leishmania* infection was detected by serology or PCR in dogs in seven of the 11 districts where human VL cases had been notified, and ITS1 sequences identical to *L. infantum* reference sequences were obtained from two dogs. These data provide a documented baseline of canine infection in these communities. As Albania has no national canine surveillance programme, a systematic, population-based canine survey would be needed to estimate the current frequency and distribution of infection and to assess its relationship with human VL.

## Acknowledgements

The authors thank the regional veterinary services for their assistance with field sampling.

## Ethical approval

Blood samples were collected by official veterinarians of the regional veterinary authorities as part of the investigation of notified human VL cases, with the consent of the dog owners.

## Author Contributions

Conceptualization: TM, GB; Methodology: TM, KM, SB; Formal analysis: TM, YNB; Visualization: KM; Writing – original draft: TM, KM, GB; Writing – review and editing: TM, KM, YNB, SB, GB. All authors have read and agreed to the published version of the manuscript.

## Data Availability Statement

The data that support the findings of this study belong to the national surveillance infrastructure and are not publicly accessible. Data are, however, available from the Institute of Public Health upon reasonable request.

## Funding data

This research received no external funding.

## References

Adhami, J., & Murati, N. (1977). Canine leishmaniasis and the natural reservoir of visceral leishmaniasis in Albania. Buletini i Universitetit të Tiranës, Seria Shkencat Mjeksore, 4, 69–74. (In Albanian)

Ayhan, N., Velo, E., de Lamballerie, X., Kota, M., Kadriaj, P., Özbel, Y., Charrel, R. N., & Bino, S. (2016). Detection of Leishmania infantum and a novel phlebovirus (Balkan virus) from sand flies in Albania. Vector-Borne and Zoonotic Diseases, 16(12), 802–806. 10.1089/vbz.2016.2002

Baneth, G., Dank, G., Keren-Kornblatt, E., Sekeles, E., Adini, I., Eisenberger, C. L., Schnur, L. F., King, R., & Jaffe, C. L. (1998). Emergence of visceral leishmaniasis in central Israel. The American Journal of Tropical Medicine and Hygiene, 59(5), 722–725. 10.4269/ajtmh.1998.59.722

Baneth, G., Koutinas, A. F., Solano-Gallego, L., Bourdeau, P., & Ferrer, L. (2008). Canine leishmaniosis – new concepts and insights on an expanding zoonosis: Part one. Trends in Parasitology, 24(7), 324–330. 10.1016/j.pt.2008.04.001

Baneth, G., Nachum-Biala, Y., Zuberi, A., Zipori-Barki, N., Orshan, L., Kleinerman, G., Shmueli-Goldin, A., Bellaiche, M., Leszkowicz-Mazuz, M., Salant, H., & Yasur-Landau, D. (2020). Leishmania infection in cats and dogs housed together in an animal shelter reveals a higher parasite load in infected dogs despite a greater seroprevalence among cats. Parasites & Vectors, 13(1), 115. 10.1186/s13071-020-3989-3

Bino, S., Vasili, A., Myrseli, T., Dikolli, E., Mersini, K., Alla, L., Sulo, J., Crilly, J., Yotebieng, K., & Bersani, A. M. (2016). One Health approach gap analysis of leishmaniasis detection, prevention, and response in the Republic of Albania [Abstract]. International Journal of Infectious Diseases, 53(Suppl.), 151.

Cani, E., Myrseli, T., Petrela, R., Minarolli, P., & Pano, K. (2001). Visceral leishmaniasis – a zoonosis with high potential risk in Albania. Revista Veterinaria, 5, 81–92. (In Albanian)

Cani, E., Petrela, R., Myrseli, T., Fuga, L., Minarolli, P., & Hyskaj, M. (2001). Identification of two visceral Leishmania strains isolated in Tirana, Albania [Abstract]. Clinical Microbiology and Infection, 7(Suppl. 1), 346.

Carvalho, B. M., Maia, C., Courtenay, O., Llabrés-Brustenga, A., Lotto Batista, M., Moirano, G., van Daalen, K. R., Semenza, J. C., & Lowe, R. (2024). A climatic suitability indicator to support Leishmania infantum surveillance in Europe: A modelling study. The Lancet Regional Health – Europe, 43, 100971. 10.1016/j.lanepe.2024.100971

Cicko, Z., Zanaj, S., Kusi, I., & Cani, E. (1999). Kërkime mbi leishmaniozën kanine në Shqipëri [Research on canine leishmaniosis in Albania]. Buletini i Shkencave Bujqësore, 3, 109–113.

EFSA Panel on Animal Health and Welfare. (2015). Scientific opinion on canine leishmaniosis. EFSA Journal, 13(4), Article 4075. 10.2903/j.efsa.2015.4075

Francino, O., Altet, L., Sánchez-Robert, E., Rodriguez, A., Solano-Gallego, L., Alberola, J., Ferrer, L., Sánchez, A., & Roura, X. (2006). Advantages of real-time PCR assay for diagnosis and monitoring of canine leishmaniosis. Veterinary Parasitology, 137(3–4), 214–221. 10.1016/j.vetpar.2006.01.011

Hidé, M., Michel, G., Legueult, K., Pin, R., Leonard, S., Simon, L., Bañuls, A.-L., Delaunay, P., Marty, P., & Pomares, C. (2024). Asymptomatic Leishmania infantum infection in dogs and dog owners in an endemic area in southeast France. Parasite, 31, 16. 10.1051/parasite/2024019

Kero, A., & Xinxo, A. (1998). Epidemiological characteristics of leishmaniasis in Albania in the period 1984–1996. Giornale Italiano di Medicina Tropicale, 3, 55–57.

Laidoudi, Y., Sevestre, J., Bedjaoui, S., Watier-Grillot, S., & Davoust, B. (2024). Serosurvey of canine leishmaniasis in five departments near an identified human clinical case in Marseille (France). One Health, 19, 100855. 10.1016/j.onehlt.2024.100855

Lazri, T., Duscher, G., Edelhofer, R., Bytyci, B., Gjino, P., & Joachim, A. (2008). Infektionen mit arthropodenübertragenen Parasiten bei Hunden im Kosovo und in Albanien unter besonderer Berücksichtigung der Leishmanieninfektionen [Arthropod-borne parasites of dogs, especially Leishmania, in Kosovo and Albania]. Wiener Klinische Wochenschrift, 120(Suppl. 4), 54–58. 10.1007/s00508-008-1076-4

Maia, C., Conceição, C., Pereira, A., Rocha, R., Ortuño, M., Muñoz, C., Jumakanova, Z., Pérez-Cutillas, P., Özbel, Y., Töz, S., Baneth, G., Monge-Maillo, B., Gasimov, E., Van der Stede, Y., Torres, G., Gossner, C. M., & Berriatua, E. (2023). The estimated distribution of autochthonous leishmaniasis by Leishmania infantum in Europe in 2005–2020. PLoS Neglected Tropical Diseases, 17(7), e0011497. 10.1371/journal.pntd.0011497

Miho, V., Adhami, J., Hobdari, G., & Murati, N. (1989). Observation concerning a visceral leishmaniasis focal point. Revista Mjekësore, 5, 13–15. (In Albanian)

Morales-Yuste, M., Martín-Sánchez, J., & Corpas-López, V. (2022). Canine leishmaniasis: Update on epidemiology, diagnosis, treatment, and prevention. Veterinary Sciences, 9(8), 387. 10.3390/vetsci9080387

Myrseli, T., Mersini, K., Kondi, E., & Allmuca, H. (2009). Një studim seroepidemiologjik për leishmaniazën viscerale kanine në rrethin e Tiranës gjatë periudhës 2007–2008 [A seroepidemiological study of canine visceral leishmaniasis in the district of Tirana during 2007–2008]. Revista Mjekësore, 4, 74–82.

Myrseli, T., Rogozi, E., & Simaku, A. (2021). Të dhënat e para të leishmaniazës viscerale asimptomatike në njerëz në Shqipëri: Një studim retrospektiv [First data on asymptomatic human visceral leishmaniasis in Albania: A retrospective study]. Buletini i Institutit të Shëndetit Publik, 4, 29–34.

Myrseli, T., Simaku, A., Baci, M., & Bino, S. (2014). Leishmanioza viscerale dhe kutane në Shqipëri: Analizë retrospektive për 2005–2013 [Visceral and cutaneous leishmaniasis in Albania: A retrospective analysis for 2005–2013]. Buletini i Institutit të Shëndetit Publik, 4, 4–18.

Myrseli, T., Velo, E., Bino, S., Alla, L., Çomo, N., Mersini, K., Gradoni, L., & Maroli, M. (2011, April 25–30). Visceral and cutaneous leishmaniasis in Albania [Abstract]. In Proceedings of the 7th International Symposium on Phlebotomine Sandflies (p. 143). Kuşadasi, Turkey.

Symeonidou, I., Angelou, A., Theodoridis, A., Sioutas, G., & Papadopoulos, E. (2021). Canine leishmaniosis in Greece: An updated countrywide serological study and associated risk factors. Pathogens, 10(9), 1129. 10.3390/pathogens10091129

Talmi-Frank, D., Nasereddin, A., Schnur, L. F., Schönian, G., Töz, S. Ö., Jaffe, C. L., & Baneth, G. (2010). Detection and identification of Old World Leishmania by high resolution melt analysis. PLoS Neglected Tropical Diseases, 4(1), e581. 10.1371/journal.pntd.0000581

Travi, B. L., Cordeiro-da-Silva, A., Dantas-Torres, F., & Miró, G. (2018). Canine visceral leishmaniasis: Diagnosis and management of the reservoir living among us. PLoS Neglected Tropical Diseases, 12(1), e0006082. 10.1371/journal.pntd.0006082

Tzani, M., Barrasa, A., Vakali, A., Georgakopoulou, T., Mellou, K., & Pervanidou, D. (2021). Surveillance data for human leishmaniasis indicate the need for a sustainable action plan for its management and control, Greece, 2004 to 2018. Eurosurveillance, 26(18), 2000159. 10.2807/1560-7917.ES.2021.26.18.2000159

Vaselek, S. (2021). Systematic review: Re-emergence of human leishmaniasis in the Balkans. Tropical Medicine & International Health, 26(10), 1189–1199. 10.1111/tmi.13653

Velo, E., Bongiorno, G., Kadriaj, P., Myrseli, T., Crilly, J., Lika, A., Mersini, K., Di Muccio, T., Bino, S., Gramiccia, M., Gradoni, L., & Maroli, M. (2017). The current status of phlebotomine sand flies in Albania and incrimination of Phlebotomus neglectus (Diptera, Psychodidae) as the main vector of Leishmania infantum. PLoS ONE, 12(6), e0179118. 10.1371/journal.pone.0179118

Xhekaj, B., Hoxha, I., Platzgummer, K., Kniha, E., Walochnik, J., Sherifi, K., Rexhepi, A., Behluli, B., Dvořák, V., Fuehrer, H.-P., Obwaller, A. G., Poeppl, W., Stefanovska, J., & Cvetkovikj, A. (2023). First detection and molecular analysis of Leishmania infantum DNA in sand flies of Kosovo. Pathogens, 12(10), 1190. 10.3390/pathogens12101190

