## Supplementary File for "Canine *Leishmania* infection in communities with notified human visceral leishmaniasis cases in Albania"

**Supplementary File 1. ITS1 sequences and GenBank comparison of *Leishmania* detected in two dogs (Albania, 2015)**

**S1. Methods summary**

Whole-blood samples positive by kinetoplast DNA (kDNA) real-time PCR were analysed by internal transcribed spacer 1 (ITS1) high-resolution melt PCR with primers ITS1-F-219F/ITS1-R-219R (Talmi-Frank et al., 2010). Amplicons from the two ITS1-positive samples were sequenced and the trimmed sequences were compared with the GenBank nucleotide collection using BLASTn.

**S2. ITS1 sequences (FASTA)**

The two sequences were obtained from the two ITS1-positive dogs, from Fier and Lezha districts.

*>Sample_A_ITS1*
ATTACACCCAAAAAACATATACAACTCGGGGAGACCTATGTATATATATGTAGGCCT
TTCCCACATACACAGCAAAGTTTTGTACTCAAAATTTGCAGTAAAAAAAAGGCCG
ATCGACGTTATAACGCACCGCCTATACAAAAGCAAAAATGTCCGTTTATACAAAAA
ATATACGGCGTTTCGGTTTTTGGCGGGGTGGGTGCGTGTGTGGATAACGG

*>Sample_B_ITS1*
ATTACACCCAAAAAACATATACAACTCGGGGAGACCTATGTATATATATGTAGGCCT
TTCCCACATACACAGCAAAGTTTTGTACTCAAAATTTGCAGTAAAAAAAAGGCCG
ATCGACGTTATAACGCACCGCCTATACAAAAGCAAAAATGTCCGTTTATACAAAAA
ATATACGGCGTTTCGGTTTTTGGCGGGGTGGGTGCGTGTGTGGATAACGG

Both sequences are 218 bp long and identical to each other.

**S3. Closest GenBank matches**

| Query | Length (bp) | Closest GenBank matches (identical to each other) | Query coverage | Identity | Accession numbers |
|---|---|---|---|---|---|
| Sample_A_ITS1 | 218 | *L. infantum* isolates MHOM/CN/2009/KSF-4, KSF-6 and KSF-7 | 100% | 100% (218/218) | KU975157.1<br>KU975158.1<br>KU975159.1 |
| Sample_B_ITS1 | 218 | *L. infantum* isolates MHOM/CN/2009/KSF-4, KSF-6 and KSF-7 | 100% | 100% (218/218) | KU975157.1<br>KU975158.1<br>KU975159.1 |
